# Humanin analogue promotes metabolic reprogramming to protect the ischemic heart

**DOI:** 10.64898/2026.06.16.732776

**Authors:** Zhenwei Gong, Ebin Johny, Ye Liu, Sivakama Bharathi, Satish Vasamsetti, Eric Goetzman, Partha Dutta, Radhika Muzumdar

## Abstract

**Background:** Myocardial ischemia drives adverse cardiac remodeling, metabolic inflexibility, and progression to heart failure. Mitochondrial dysfunction and impaired substrate utilization contribute to cardiomyocyte death and fibrosis, particularly with aging. Humanin (HNG), a mitochondria-derived peptide, has been shown to reduce acute ischemic injury, but its role in chronic ischemia and cardiac remodeling remains unknown.

**Methods:** We investigated the effects of HNG treatment in young and aged murine models of myocardial ischemia without reperfusion. Cardiac function and structure were assessed by echocardiography and molecular markers of remodeling. Myocardial metabolism was interrogated using targeted metabolomics, gene expression, substrate uptake assays, and metabolic flux analyses. Mechanistic studies examined glucose transporter trafficking and protein–protein interactions.

**Results:** HNG treatment improved cardiac function and significantly attenuated adverse remodeling in both young and old mice. HNG treatment induced marked metabolic reprogramming characterized by reduced myocardial fatty acid content, downregulation of fatty acid uptake and oxidation pathways, and decreased oxidative stress. Importantly, these changes were accompanied by enhanced glucose oxidation, increased tricarboxylic acid cycle flux, improved coupling of glycolysis to mitochondrial oxidation, and increased ATP production. Time-course studies demonstrated that increased glucose oxidation preceded reductions in fatty acid oxidation, indicating a primary role for glucose metabolism in HNG-mediated cardioprotection. Mechanistically, we identified vesicle-associated membrane protein 7 (VAMP7) as a novel binding partner of HNG, and that this interaction is required for GLUT4 translocation to the plasma membrane and HNG-induced ATP generation.

**Conclusions:** HNG protects the ischemic heart by promoting metabolic reprogramming that shifts substrate utilization from fatty acids to glucose and limiting maladaptive remodeling. These findings identify HNG as a novel regulator of cardiac metabolism and a potential therapeutic strategy for ischemic heart failure.

**Graphical abstract:** 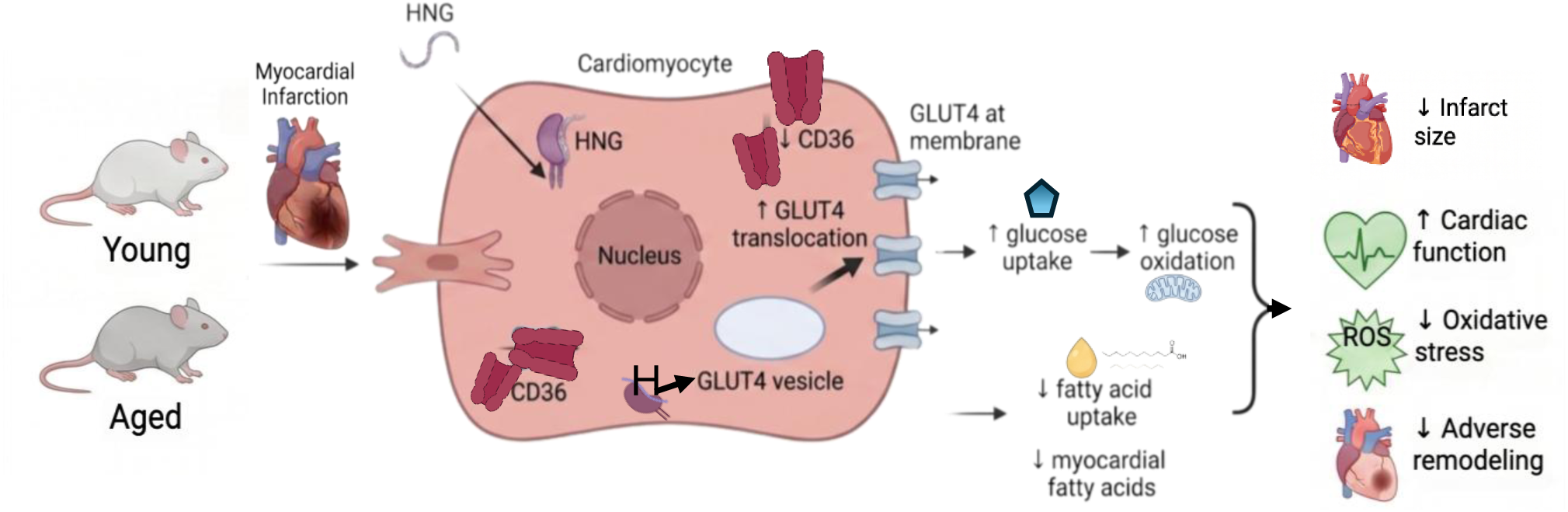

**What are the clinical implications?:** Heart failure (HF) is a major global health concern, affecting over 6.7 million adults in the United States alone, with projections to exceed 11 million by 2050. Myocardial infarction (MI) is a leading cause of HF. Despite substantial advances in acute MI care, survivors remain at high risk for adverse cardiac remodeling and chronic HF, especially in the elderly. We report here that treatment with a potent analog of Humanin (HN), an endogenous mitochondria-associated peptide, decreases infarct size, decreases fibrosis and improves cardiac function following cardiac ischemia induced by permanent ligation of coronary artery in both young and aged mice. These effects are associated with changes in cardiac metabolism, oxidative stress, and remodeling. HN and analogs have been shown to be beneficial in many age-related diseases. The endogenous origin of Humanin, its favorable safety profile in preclinical studies and its pleiotropic effects support targeting HNG as a promising therapeutic strategy for ischemic heart disease and post–myocardial infarction heart failure in humans.

## INTRODUCTION

Heart failure (HF) is a major global health concern, affecting over 6.7 million adults in the United States alone—a number projected to exceed 11 million by 2050. With a lifetime risk of approximately 1 in 4, and accounting for nearly half of all cardiovascular-related deaths, HF represents a substantial and growing clinical and economic burden. This burden is further exacerbated by the increasing prevalence of comorbidities such as diabetes, hypertension, and chronic kidney disease (1–3) . Annual HF-related healthcare costs in the United States are projected to triple by 2050, rising to approximately $850 billion, underscoring the urgent need for improved preventive and therapeutic strategies (1). Ischemic heart disease, particularly myocardial infarction (MI), remains a leading cause of HF. Despite substantial advances in acute MI care, survivors remain at high risk for adverse cardiac remodeling, chronic HF, and long-term mortality (3). Indeed, long-term follow-up studies report 8-year mortality rates approaching 65% following MI (4). Given the magnitude of this burden, along with the heightened vulnerability of individuals over 65 years of age, there is a critical need to develop therapies that target post-MI remodeling and limit HF progression across the lifespan.

The current standard of care for acute MI emphasizes rapid reperfusion via primary angioplasty or thrombolysis to reduce infarct size and improve short-term survival. However, restoration of blood flow paradoxically induces ischemia–reperfusion (I/R) injury, exacerbating myocardial damage. Reperfusion triggers oxidative stress and calcium overload in previously ischemic cardiomyocytes, leading to mitochondrial dysfunction, opening of the mitochondrial permeability transition pore, and cell death (5). This secondary injury amplifies inflammation and fibrosis, driving adverse ventricular remodeling and HF progression. Older MI patients are particularly vulnerable due to age-related oxidative stress burden, frailty, and associated comorbidities, and may poorly tolerate aggressive interventions. Despite extensive investigation, cardioprotective strategies such as ischemic conditioning and pharmacologic approaches have shown limited success in clinical translation (5).

Therapeutic approaches targeting myocardial energy metabolism have shown promise in preclinical models of ischemic injury. Experimental studies demonstrate that promoting glucose oxidation or inhibiting fatty acid oxidation (FAO) during ischemia can reduce infarct size and improve cardiac function. Agents such as dichloroacetate, which activates pyruvate dehydrogenase, and etomoxir, an inhibitor of fatty acid β-oxidation, have demonstrated cardioprotective effects in animal models (6, 7). However, clinical translation of these agents has been limited by safety and tolerability concerns, including dose-limiting neuropathy with dichloroacetate and hepatotoxicity associated with etomoxir (7). These limitations highlight the need for novel metabolic interventions that safely attenuate post-ischemic injury and remodeling, particularly in the growing population of older MI patients at the highest risk for HF.

Humanin (HN), a mitochondria-derived peptide, and its analogs have emerged as promising therapeutic candidates at the intersection of cardiovascular metabolism and aging. HN is a 24–amino acid peptide encoded within the 16S rRNA region of mitochondrial DNA and was first identified for its neuroprotective effects in models of Alzheimer’s disease (8). HN is expressed in multiple tissues, including the heart, skeletal muscle, and central nervous system. HN and its analogs have demonstrated protective efficacy in multiple age-associated diseases, including stroke, atherosclerosis, and myocardial ischemia, without evidence of toxicity in preclinical models (9, 10). Our laboratory has investigated the HN analog, Gly¹⁴Humanin (HNG) in the context of myocardial I/R injury, with encouraging results in both rodent and porcine models (11, 12).

In the present study, we examined the effects of daily HNG treatment for 28 days on progression to heart failure following chronic myocardial ischemia in young and old mice. We assessed cardiac contractile performance and indices of remodeling and interrogated underlying mechanisms using unbiased myocardial metabolomics, complemented by targeted in vivo and in vitro analyses of fatty acid and glucose uptake and oxidation.

## MATERIALS AND METHODS

### Animals

Young male and female C57BL/6J mice (3 months old) were purchased from The Jackson Laboratory and housed in individually ventilated cages at the UPMC Animal Facility. Aged mice (20–24 months old) were obtained from the National Institute on Aging (NIA). All animals received veterinary care under the supervision of the Division of Laboratory Animal Resources (DLAR) at the University of Pittsburgh. All procedures were conducted in accordance with National Institutes of Health (NIH) guidelines and approved by the Institutional Animal Care and Use Committee (IACUC) of the University of Pittsburgh. The facility is accredited by the Association for Assessment and Accreditation of Laboratory Animal Care International (AAALAC).

### Myocardial Infarction (MI)

MI was induced as previously described (13). Briefly, mice were anesthetized with 3% isoflurane delivered in oxygen (1 L/min), intubated, and mechanically ventilated at 137 breaths/min with a tidal volume of 0.18 mL. Following a left thoracotomy at the third intercostal space, the left coronary artery (LCA) was permanently ligated using an 80-nylon suture. Successful ligation was confirmed by the blanching of the myocardium. Postoperative care included thermal support, continuous monitoring until full recovery, and administration of extended-release buprenorphine (3.25 mg/kg, subcutaneous). Following MI, mice were randomized to receive either scrambled peptide (SP) or the potent humanin analog Gly¹⁴Humanin (HNG; 2 mg/kg). Treatment was initiated 24 hours post-MI via intravenous injection and continued daily via intraperitoneal administration for 28 days. A schematic representation of the study protocol is shown in Fig 1.

**Fig. 1:**
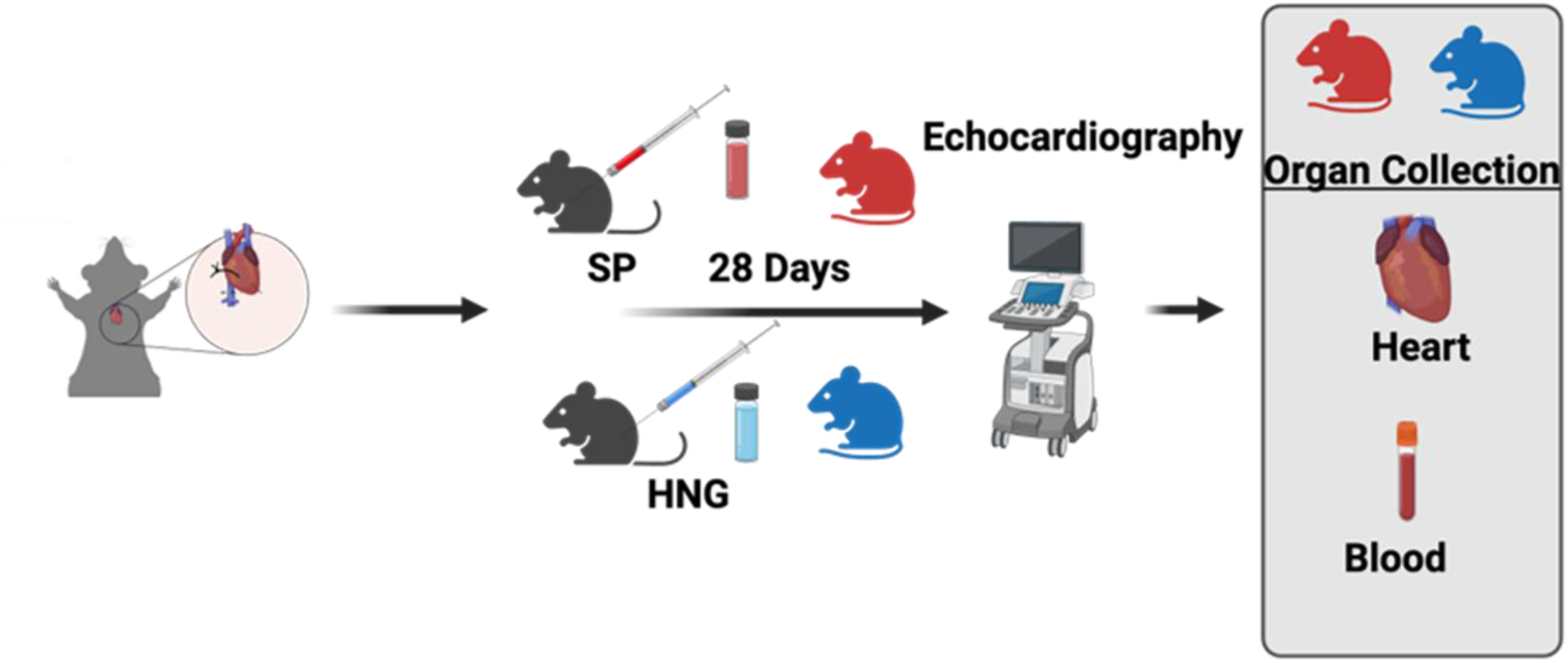
Schematic figure of experimental design.

### Echocardiography

Transthoracic echocardiography was performed using a Vevo 3100 preclinical imaging platform to assess left ventricular function. Parameters including ejection fraction, stroke volume, cardiac output, end systolic volume, and end diastolic volume were quantified from B-mode images acquired in the parasternal long axis view using VevoLAB software (version 3.2.6).M-mode images were acquired at the level of the papillary muscles to visualize left ventricular wall motion.

### In vivo fatty acid uptake assay

One hour before tissue harvest after MI and 28 days of SP or HNG treatment, a group of young female mice received a single dose (0.5mg/kg) injection of BODIPY labeled palmitate (Life Technologies, Cat# D3821). Heart tissue was collected and fixed for examination of green fluorescence.

### Organ Harvesting

Mice were euthanized in accordance with University of Pittsburgh IACUC guidelines. Peripheral blood was collected by cardiac puncture or retroorbital sinus sampling, and serum was isolated by centrifugation. Following perfusion with phosphate buffered saline (PBS) via the left ventricle, hearts were harvested and processed for histology, immunofluorescence staining, immunoblotting, or quantitative PCR (qPCR).

### Histology and Immunofluorescence

#### Masson’s Trichrome Staining

Ventricular heart tissue was embedded in optimal cutting temperature (OCT) compound (Fisher Healthcare) on dry ice. Cryosections (10 µm) were prepared at the infarct level. Fibrosis was assessed using Masson’s trichrome staining (StatLab™ MasterTech Trichrome Kit, Cat# STLKMTRPT). Sections were fixed overnight in Bouin’s solution, rinsed thoroughly, stained with hematoxylin A/B (1:1), scarlet acid fuchsin, phosphomolybdic acid, and aniline blue, followed by dehydration and mounting with Permount (Fisher SP15-100). Imaging was performed using an Evident Scientific VS200 slide scanner.

#### Cardiomyocyte Size

Cardiomyocyte hypertrophy was assessed by plasma membrane staining using wheat germ agglutinin (WGA). Sections (8 µm) were incubated with Alexa Fluor 594-conjugated WGA (Thermo Fisher, W11262) and mounted with DAPI containing fluorescence medium (Southern Biotech, OB010020). Images were acquired on a Nikon A1 spectral confocal microscope, and cardiomyocyte area was quantified using ImageJ.

#### Immunofluorescence

Sections were fixed in 4% paraformaldehyde, permeabilized with 0.1% Triton X100, and incubated with primary antibodies against CD68 (Invitrogen, 14068182) and 8-OHdG (Novus Bio, NB6001508). Secondary antibodies included donkey anti-goat AF595 (Invitrogen, A11058) and donkey anti-goat AF647 (Invitrogen, A21447). Nuclei were counterstained with DAPI Fluoromount (Southern Biotech, 010020). Imaging and analysis were performed using a Nikon A1 confocal microscope and ImageJ.

#### Serum B-type natriuretic peptide (BNP)

Serum BNP levels were measured using a mouse BNP ELISA kit (MyBioSource, Cat# MBS2510603) according to the manufacturer’s instructions.

#### Cell Culture

H9C2 cells (ATCC, CRL1446) were cultured in DMEM containing 5 mM D-glucose, 1 mM sodium pyruvate, and 10% FBS (Biowest) at 37°C in 10% CO₂. GFP and GFP-HN expressing stable lines were generated using lentiviral vectors and sorted by flow cytometry. HEK293 cells (ATCC) were cultured in DMEM with 25 mM D-glucose, 1 mM sodium pyruvate, and 10% FBS and transfected using Lipofectamine 3000 (Thermo Fisher). HL1 cardiomyocytes (Sigma Millipore) were cultured in Claycomb medium supplemented with 10% FBS, 0.1 mM norepinephrine, 2 mM L-glutamine, and 1% penicillin/streptomycin.

#### Co-Immunoprecipitation and Proteomics

GFP and GFP-HN expressing H9C2 cells were lysed in IP buffer (150 mM NaCl, 50 mM Tris HCl pH 8.0, 1% Triton X100) with protease and phosphatase inhibitors. Lysates were incubated overnight with mouse anti-GFP antibody, followed by protein G bead incubation for 2 hours. After washing, proteins were eluted in buffer containing 50 mM TEAB and 5% SDS. Proteomics was performed in Health Sciences Mass Spectrometry Core at the University of Pittsburgh.

#### Immunoblotting

Protein lysates were prepared in RIPA buffer with protease and phosphatase inhibitors. Protein concentrations were determined by BCA assay. Thirty micrograms of protein were resolved by SDS-PAGE, transferred to nitrocellulose membranes, blocked with nonfat milk, and incubated with primary antibodies overnight. Detection was performed using HRP-conjugated secondary antibodies and chemiluminescence (Thermo Fisher) on a LAS3000 imaging system (Fujifilm). Densitometry was performed using ImageJ.

#### Metabolomic Profiling

Periinfarct myocardial tissues were isolated, snap frozen in liquid nitrogen, and analyzed by the West Coast Metabolomics Center at the University of California, Davis. In brief, metabolites were extracted using a modified MTBE-based liquid–liquid extraction, and the methanol/water phase containing polar metabolites was collected for analysis.Metabolomic profiling was performed by HILIC–ESI–QTOF MS/MS using an Agilent 1290 UHPLC coupled to a high-resolution QTOF instrument, with separation on a BEH amide column under a water/acetonitrile gradient containing ammonium formate and formic acid. Data were acquired in both positive and negative ionization modes, with MS/MS fragmentation for metabolite identification. Raw data were processed using MS-DIAL for peak detection, alignment, and annotation based on in-house and public spectral libraries (NIST, MoNA), followed by manual curation. Metabolite abundances were quantified using peak heights and normalized to internal standards (iTIC) to correct for instrument variability, yielding relative semi-quantitative measurements.

#### Fatty Acid Uptake Assay

Fatty acid uptake in H9C2 cells was measured using the Fatty Acid Uptake Assay Kit (Abcam) following the manufacturer’s protocol. This assay is based on the uptake of a fluorescent long-chain fatty acid analog by live cells via endogenous fatty acid transporters, followed by intracellular accumulation in lipid compartments. A membrane-impermeable quenching reagent was used to eliminate extracellular fluorescence, enabling specific detection of intracellular fatty acid uptake without wash steps. H9C2 were pretreated with 1 μM of HNG for 1 hour followed by adding fluorescent probe for additional 1 hour or 24 hours. Fluorescence intensity (Ex/Em ∼485–515 nm) was quantified using a microplate reader, providing a sensitive and quantitative measure of fatty acid uptake kinetics.

#### Glucose uptake

Cellular glucose uptake was measured using a commercially available Glucose Uptake Assay Kit (Abcam) according to the manufacturer’s instructions. uptake of by cells via glucose transporters, followed by intracellular accumulation due to incomplete metabolism. H9C2 cells were plated in 96-well plates and starved for 2 hours. Cells were then treated with 1mM 2-deoxyglucose and 1 μM of HNG for 15, 30 and 60 min. Fluorescence was measured using a microplate reader at Ex/Em = ∼535/587 nm.

#### Seahorse extracellular flux analysis

H9C2 cells were counted and seeded at 10,000 cells per well onto a 96-well Seahorse plate. The following morning the media was changed to DMEM containing 10 mM glucose ± HNG peptide (1 µM) and the cells cultured for 24 hr. Then, the media was replaced with Seahorse assay media and the cells were assayed for oxygen consumption rate (OCR) with the Seahorse Extracellular Flux Analyzer. OCR values were normalized to protein concentration.

#### 14C glucose oxidation

H9C2 cells were grown in DMEM containing 10 mM glucose and treated ± HNG peptide (1 µM) for 24 hr. Each reaction contained 100,000 cells. Uniformly labeled ^14^C-glucose was added to a final concentration of 1 µCi/ml and the reactions allowed to proceed for 3 hr at 37°C. The reactions were acidified with perchloric acid to release ^14^C-CO_2_, which was captured on filter papers soaked in 1N KOH. The papers were subjected to scintillation counting, and data expressed as nmoles of ^14^C-glucose oxidized to ^14^C-CO_2_ per hour per 100,000 cells.

#### 13C glucose flux

H9C2 cells were pre-treated with or without 1 μM of HNG for 7 days. Cells were then starved for 2 hours followed by treatment with 10mM ^13^C-labeled glucose for 2 hours. A duplicate set of reactions contained no label, to allow data correction for the natural background abundance of ^13^C. Cells were snap frozen and harvested in 80% methanol. Samples were processed, extracted, and quantified at the Health Sciences Mass Spectrometry Core at the University of Pittsburgh. For ease of presentation, all 13C-enriched isotopologues were summed together (i.e., the sum of M+1, M+2, M+3, etc) and expressed as the total atomic percent enrichment (APE%) after correcting for the natural abundance of ^13^C.

#### GLUT4 Translocation Assay

HL1 cells grown in 6 well plates or on coverslips were transfected with HA-GLUT4-GFP (14) (gift from Dr. Timothy E. McGraw, Weill Cornell Medical College) for 48 hours and serum starved for 2 hours. Cells were treated with or without 1 µM of HNG for 1 hour, fixed in 4% paraformaldehyde, and stained under nonpermeabilized conditions using mouse anti-HA antibody and Alexa Fluor 594-conjugated goat anti-mouse IgG. Cell surface GLUT4 was quantified by flow cytometry (BD Fortessa) and confocal microscopy (Zeiss).

#### VAMP7 Knockdown

VAMP7 SMARTpool siRNA (Horizon Discovery Biosciences) was transfected using Lipofectamine RNAiMAX (Thermo Fisher) according to the manufacturer’s instructions.

#### Intracellular ATP Assay

ATP levels were quantified using the ATP Determination Kit (Thermo Fisher Scientific) per manufacturer’s instructions.

#### RNA Extraction and Quantitative RealTime PCR

Total RNA was extracted from young or aged female mouse hearts using the PureLink RNA Mini Kit (Thermo Fisher). cDNA was synthesized from 500 ng RNA using iScript (Bio-Rad). qPCR was performed using TaqMan assays in 10 µL reactions. Relative expression was calculated using the 2⁻ΔΔCt method and normalized to GAPDH.

### Statistics

All data are presented as mean ± SEM. Statistical analyses were performed using GraphPad Prism 6 (GraphPad Software, San Diego, CA). Comparisons between two groups at a single time point were conducted using an unpaired, two-tailed Student’s t-test. For analyses involving multiple time points or group comparisons, a two-way repeated-measures analysis of variance (ANOVA) followed by Bonferroni post hoc testing was applied. A p value < 0.05 was considered statistically significant.

## Results

### HNG treatment improves cardiac function following myocardial infarction in young and old female mice

We previously demonstrated that HNG exerts cardioprotective effects against oxidative stress and apoptosis in I/R models (11, 15). Building on these findings, we assessed the therapeutic effects of HNG on cardiac function in a murine MI model in both young and aged mice. Transthoracic echocardiography was performed on day 28 post- MI to evaluate left ventricular function in mice treated with either HNG or SP (Fig 2A). Although end diastolic and end systolic volumes remained unchanged between the two groups in young female mice (Fig. 2B and C), stroke volume was significantly increased in the HNG treated group (16.9 ± 6.25 µL) as compared to the control group (10.7 ± 4.4 µL) (Fig. 2D). HNG treated mice (23.02 ± 11.55 %) exhibited a significantly higher left ventricular ejection fraction compared to SP treated controls (12.46 ± 5.97 %) (Fig. 2E). Similarly, cardiac output was improved in HNG treated mice (7.4 ± 2.85 ml/min) relative to placebo treated animals (5.47 ± 2.4 ml/min) (Fig. 2F). Serum BNP levels were significantly lower in the HNG treated mice compared to the controls (Fig. 2G). Overall, these results indicate that HNG has beneficial effects on cardiac function following MI in young female animals.

**Fig. 2:**
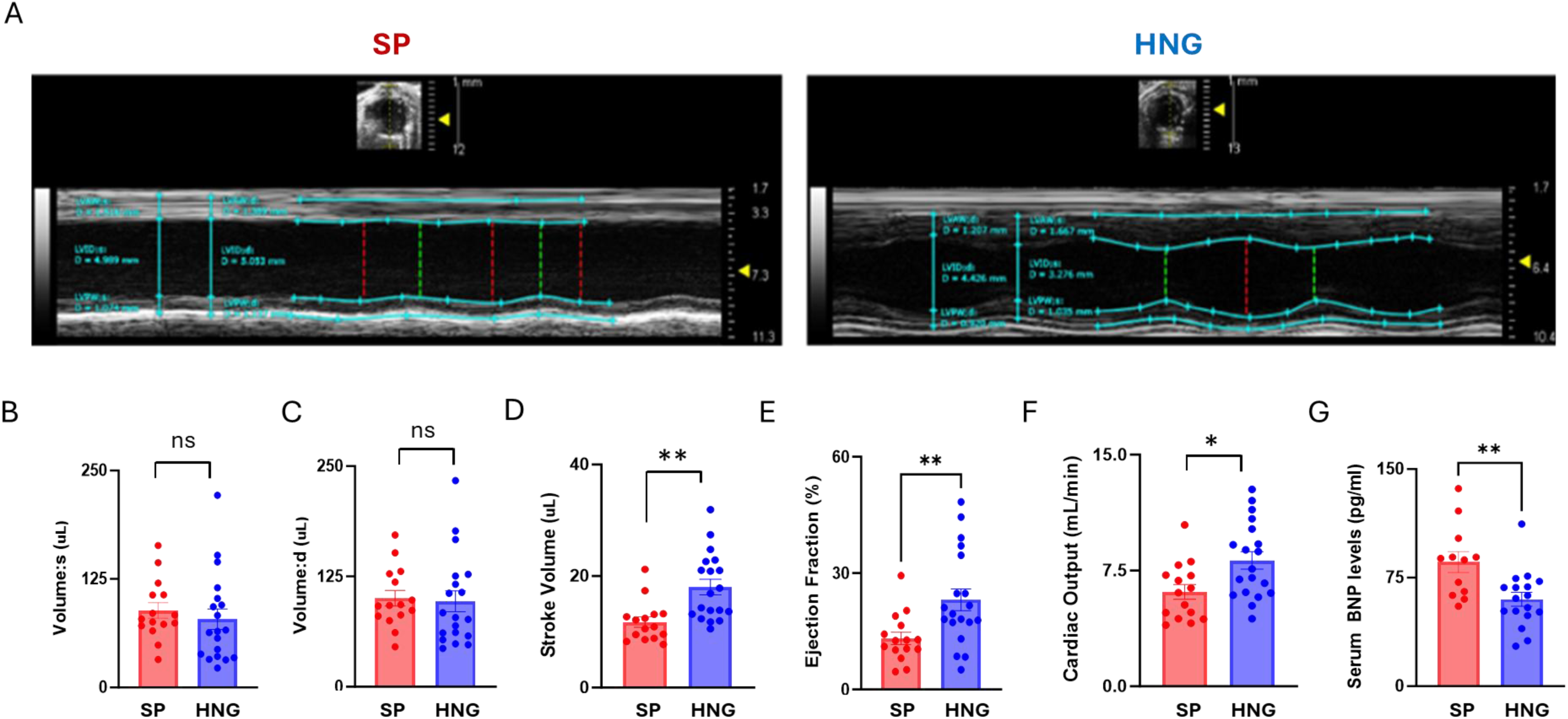
HNG improves cardiac function following myocardial ischemia in young mice. A: Representative echocardiography images of left ventricle 28 days post MI and SP or HNG treatment in young female mice; B-F: systolic volume (B), diastolic volume (C), stroke volume (D), ejection fraction (E) and cardiac output (F) in young female mice following chronic MI and SP or HNG treatment; G: serum levels of BNP in young female mice following MI and SP or HNG treatment. Results are shown as mean ± SEM. ** p<0.01

We also evaluated the cardioprotective effects of HNG in the 20-24-month-old female mice (Fig.3A) and we found similar results with unchanged end diastolic (HNG: 84.30 ± 24.05 vs Placebo: 90.16 ± 27.41 µL) and end systolic volumes (HNG: 64.57 ± 23.22 vs Placebo:76.011 ± 26.01 µL) (Fig. 3B-C) and improved stroke volume (HNG: 19.73 ± 5.24 vs Placebo: 14.15 ± 3.71µL) (Fig. 3D), ejection fraction (HNG: 24.94 ± 8.93 vs Placebo:17.19 ±7.59 %) (Fig. 3E) and cardiac output (HNG: 8.94 ± 1.81 vs Placebo: 6.20 ± 1.10 ml/min) (Fig. 3F). Serum BNP levels were significantly decreased by HNG in the old female mice (Fig. 3G).

**Fig. 3:**
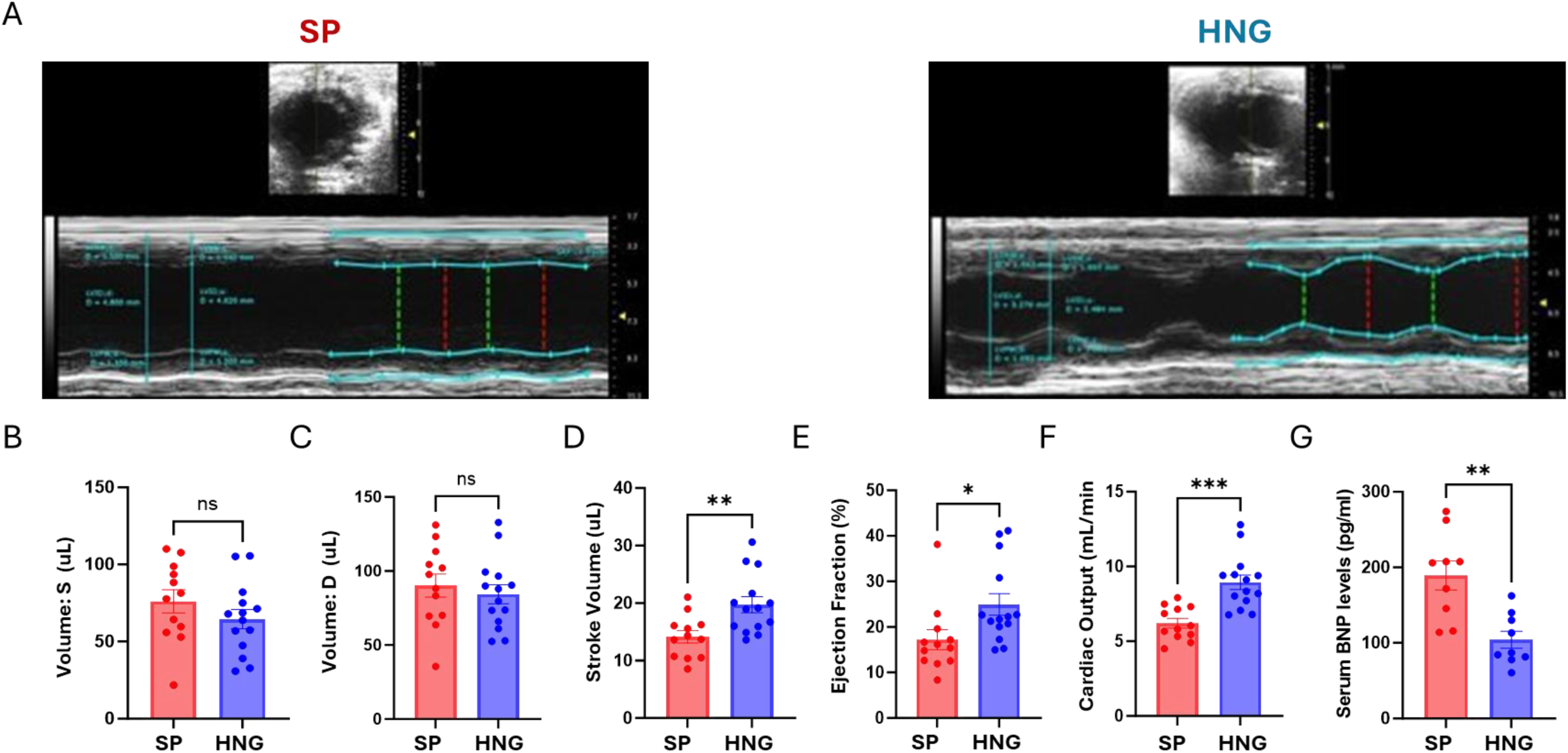
HNG improves cardiac function following myocardial ischemia in old mice. A: Representative echocardiography images of the left ventricle 28 days post MI and SP or HNG treatment in old female mice; B-F: systolic volume (B), diastolic volume (C), stroke volume (D), ejection fraction (E) and cardiac output (F) in old female mice following chronic MI and SP or HNG treatment; G: serum levels of BNP in old female mice following chronic MI and SP or HNG treatment. Results are shown as mean ± SEM. ** p<0.01

### HNG improves post-MI survival in young male mice

To assess the effect of HNG on post MI survival, MI surgery was performed in young male mice. Male mice are known to exhibit high mortality rates following MI (15), making them a relevant model for evaluating survival outcomes. Compared to SP-treated controls, HNG-treated mice demonstrated a significantly improved survival rate after MI (p<0.01) (Fig. S1).

### HNG mitigates post-MI cardiac fibrosis and cardiac hypertrophy

Histological analysis of Masson’s trichrome -stained cardiac cross-sections revealed that 28 days of HNG treatment significantly reduced cardiac collagen deposition following MI compared to SP treated controls in both young (Fig. 4A and B) and old (Fig. 4C and D) female mice (p<0.05). Quantitative PCR to examine the expression levels of fibrosis markers in the heart of both young and old female mice showed a significant decrease in the expression levels of *Tgfβ* in both young and old mice (Fig. 4E and F, p<0.05), and *Cxcl2* expression (Fig 4E, p < 0.05) in young mice with HNG treatment.

**Fig. 4:**
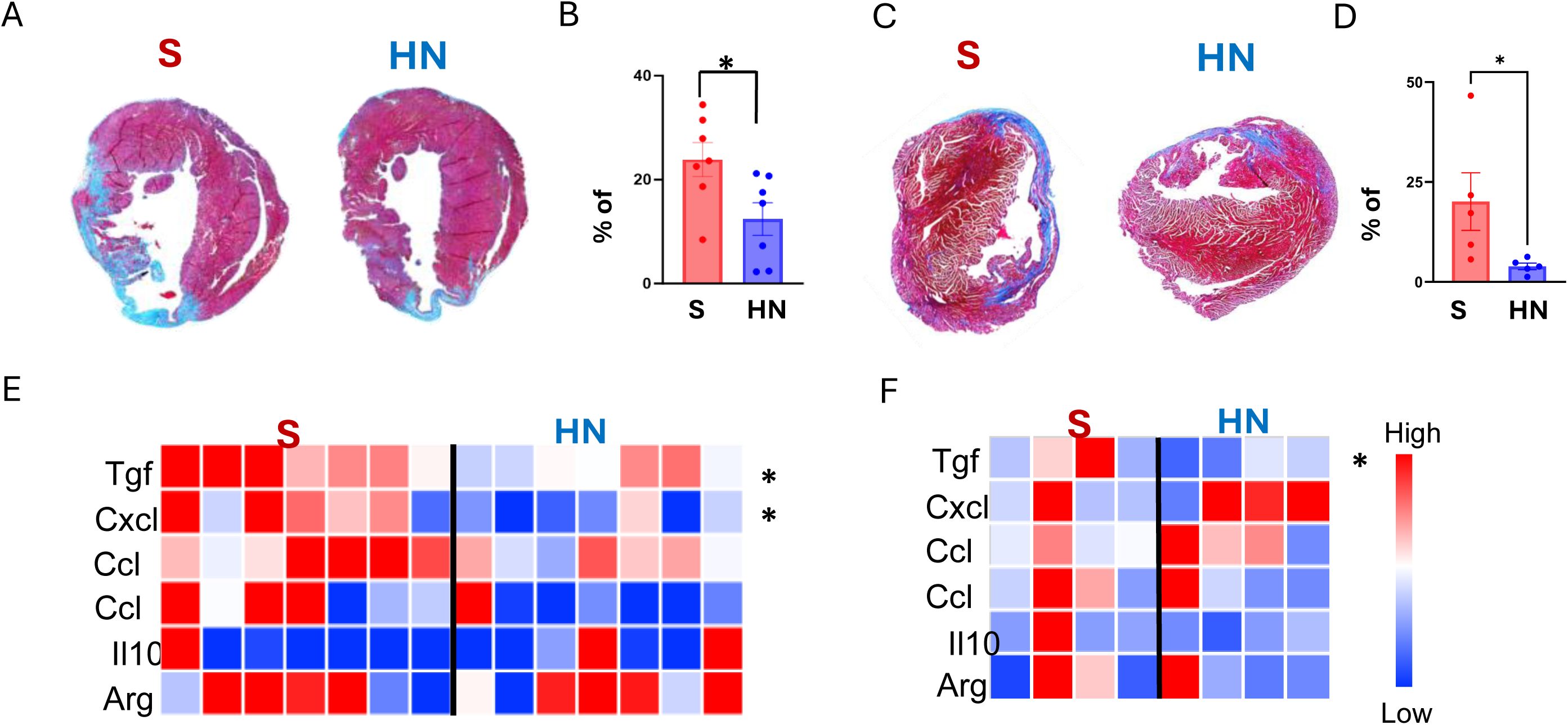
HNG reduces fibrosis following chronic MI in young and old mice. A-D: Representative cross-sectional photomicrographs of Masson’s Trichrome Staining (A and C), and quantification (B and D) in young (A and B) and old (C and D) female mice following MI and SP or HNG treatment; E-F: Relative gene expression levels of fibrosis markers in heart tissue from young (E) and old (F) female mice following chronic MI and SP or HNG treatment using quantitative real time PCR. Results are shown as mean ± SEM. * p<0.05 and ** p<0.01

In addition, Wheat Germ Agglutinin (WGA) staining demonstrated a significant reduction in cardiomyocyte size within the border zone in the HNG treated group compared to the SP treated group in both young (Fig. 5A and B, p<0.01) and old (Fig. 5C and D) female mice (P<0.001), indicating attenuation of cardiomyocyte hypertrophy. These findings suggest that HNG treatment attenuates adverse cardiac remodeling following MI by thwarting fibrosis and cardiomyocyte hypertrophy.

**Fig. 5:**
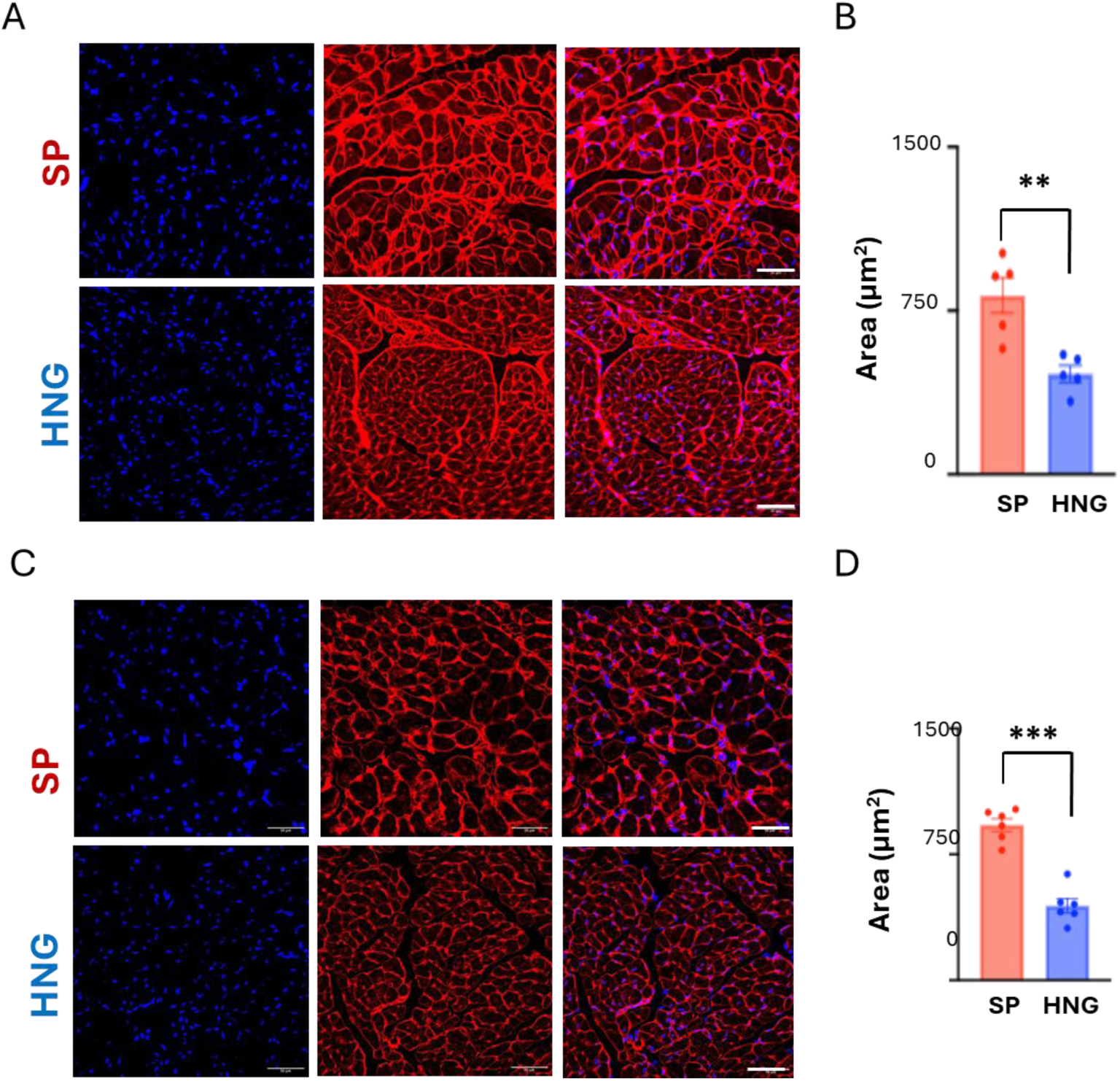
HNG treatment is associated with smaller cardiomyocyte size following chronic MI in young and old mice. A and C: Representative images for wheat germ agglutinin (WGA) staining in young (A) and old (C) female mouse hearts; B and D: quantification of the cardiomyocyte size in young (B) and old (D) female mouse hearts following MI and SP or HNG treatment. Scale bars represent 50 μm. Results are shown as mean ± SEM. ** p<0.01 and *** p<0.001

### HNG modulates oxidative DNA damage following MI

Given the observed improvements in cardiac function, reduced fibrosis, and enhanced survival with HNG treatment post-MI and the significant role of oxidative stress in cardiac biology, we assessed 8-OHdG levels as a marker of oxidative stress in the heart in both young and old female mice. Immunofluorescence imaging of cardiac tissue revealed reduced 8-OHdG levels, reflecting decreased oxidative DNA damage and attenuation of oxidative stress in young mice (Fig. 6A and B, p<0.001), while no significant changes were noted in the old mice (Fig. 6D and E). Real-time PCR results demonstrated reduced expression levels of *Nrf 2, SOD 1 and Gpx1* in young mice (Fig. 6G), while *Nrf 2* was decreased in the old mice (Fig. 6H). This finding is consistent with our previous observation that HNG exerts cardioprotective effects by reducing oxidative stress (14).

**Fig. 6:**
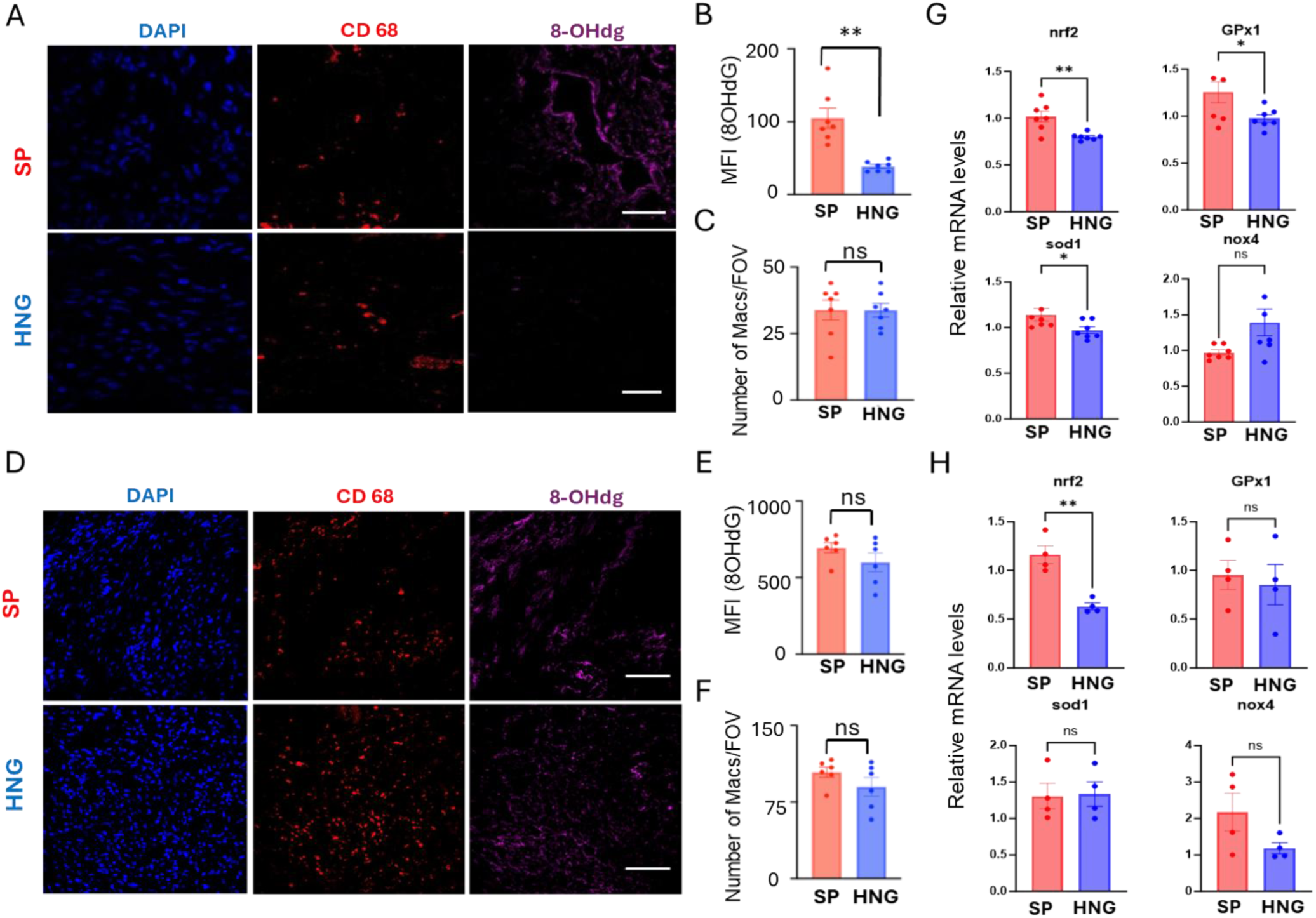
HNG treatment is associated with decreased oxidative stress following MI. A: Representative images for DAPI, CD68 and 8-OHdG staining; B-C: quantification of 8-OHdG positive staining (B) and CD68 positive macrophages (C) in young female animals treated with SP or HNG; D: Representative images for DAPI, CD68 and 8-OHdg staining; E-F: quantification of 8-OHdG-stained areas (E) and CD68 positive macrophages (F) in old female animals treated with SP or HNG; and G-H: relative mRNA expression levels for genes involved in oxidative stress, including nrf2, gpx1, sod1 and nox4 in the heart tissue in young (G) and old (H) female mice. Scale bars represent 50 μm. Results are shown as mean ± SEM. * p<0.05 and ** p<0.01

Previous studies reported that inflammatory cells are instrumental in cardiac remodeling and fibrosis. We sought to explore whether HNG suppresses MI-driven inflammation. We observed no significant difference in the number of macrophages (CD68-positive cells) between HNG and SP treated hearts in either young mice (Fig. 6A and C) or old mice (Fig. 6D and F) , suggesting that HNG’s cardioprotective effects are unlikely to be mediated by macrophages infiltrating the infarct.

### HNG treatment decreases fatty acid levels in the heart

HN and analogs have been shown to affect glucose and lipid metabolism in multiple tissues including skeletal muscle, adipose tissue, β cells, and liver (11, 16, 17).. To examine the effects of HNG treatment on cardiac metabolism, we performed untargeted metabolomics of key metabolic intermediates in the border zone of the infarct heart tissue in young female mice. Significantly reduced levels of a variety of fatty acids, including heptadecanoic acid, pentadecanoic acid, oleic acid, myristic acid, stearic acid, and linoleic acid (Fig. 7A and B) were observed. Since reduced levels of fatty acid in the heart could be a consequence of reduced uptake into cardiomyocytes and/or increased fatty acid oxidation, we examined the effects of HNG on both fatty acid uptake and oxidation. *In vivo* studies showed that HNG decreased fatty acid uptake in the heart as evidenced by BODIPY labeled palmitate uptake (Fig. 8A and B, p<0.05). This is supported by reduced mRNA expression levels of key fatty acid transport proteins such as *Fabp1* and *Cd36* in both young (Fig. 8D) and old (Fig. 8E) heart following treatment with HNG. *In vitro* studies showed that the decrease in fatty acid uptake in H9C2 cardiomyocytes was noted after 24 hours of treatment with HNG (Fig. 8C). The expression levels of key fatty acid oxidation genes were found to be decreased in the heart tissue of young (*CPT1α*, Fig. 8F, p< 0.05) and old (*CPT1α*, *PGC-1α and PPARα,* Fig. 8G, p< 0.01) mice following HNG treatment. Together, these results indicate that the lower levels of fatty acids in the heart are not due to increased fatty acid oxidation, but rather due to decreased fatty acid uptake following HNG treatment.

**Fig. 7:**
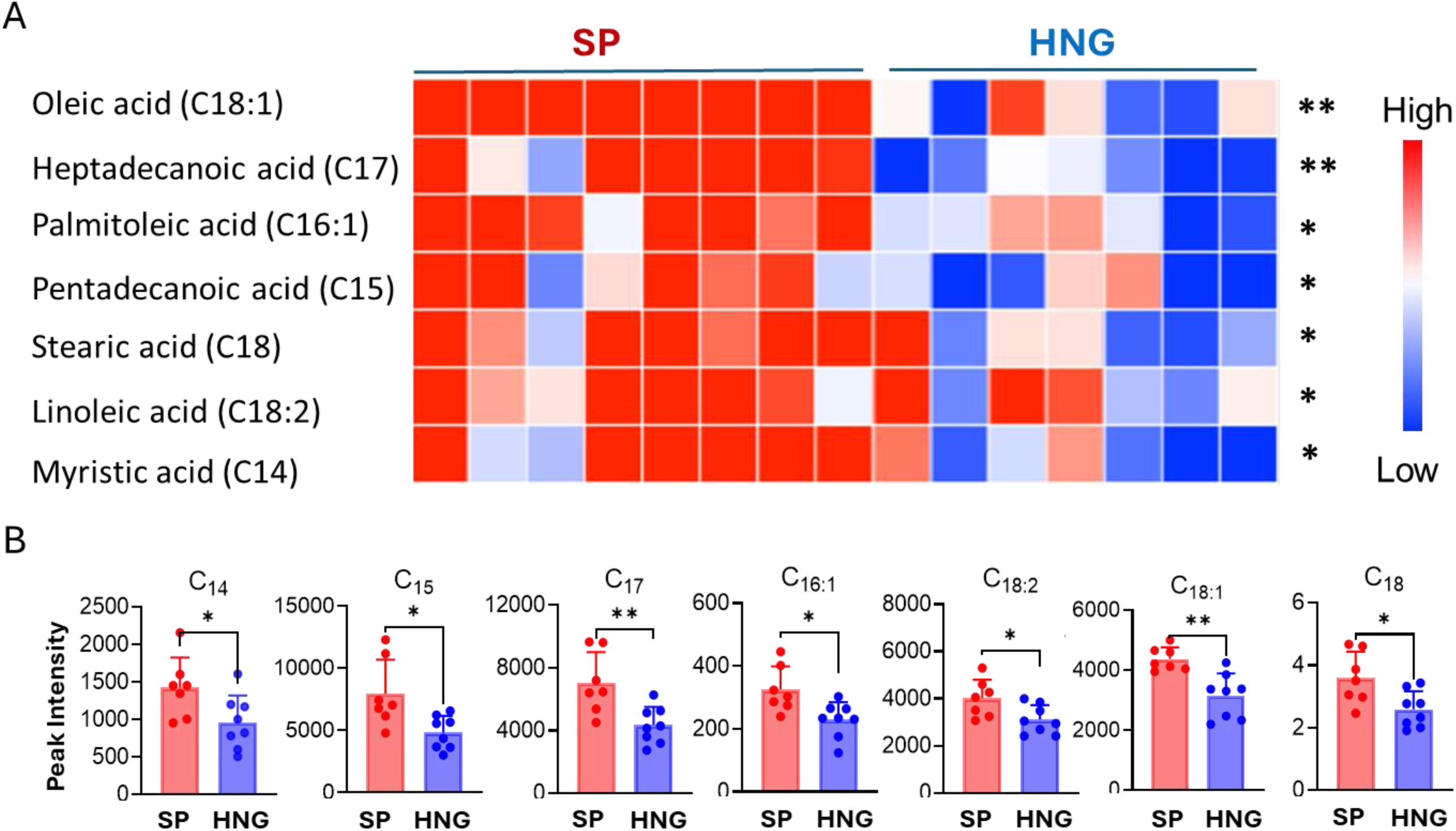
HNG treatment decreases free fatty acids in the heart following MI in young mice. A and B: heat map (A) and peak intensity (B) of free fatty acids in the border zone of the heart tissue from young female mice following MI and SP or HNG treatment in a metabolomic study. Results are shown as mean ± SEM. * p<0.05 and ** p<0.01

**Fig. 8:**
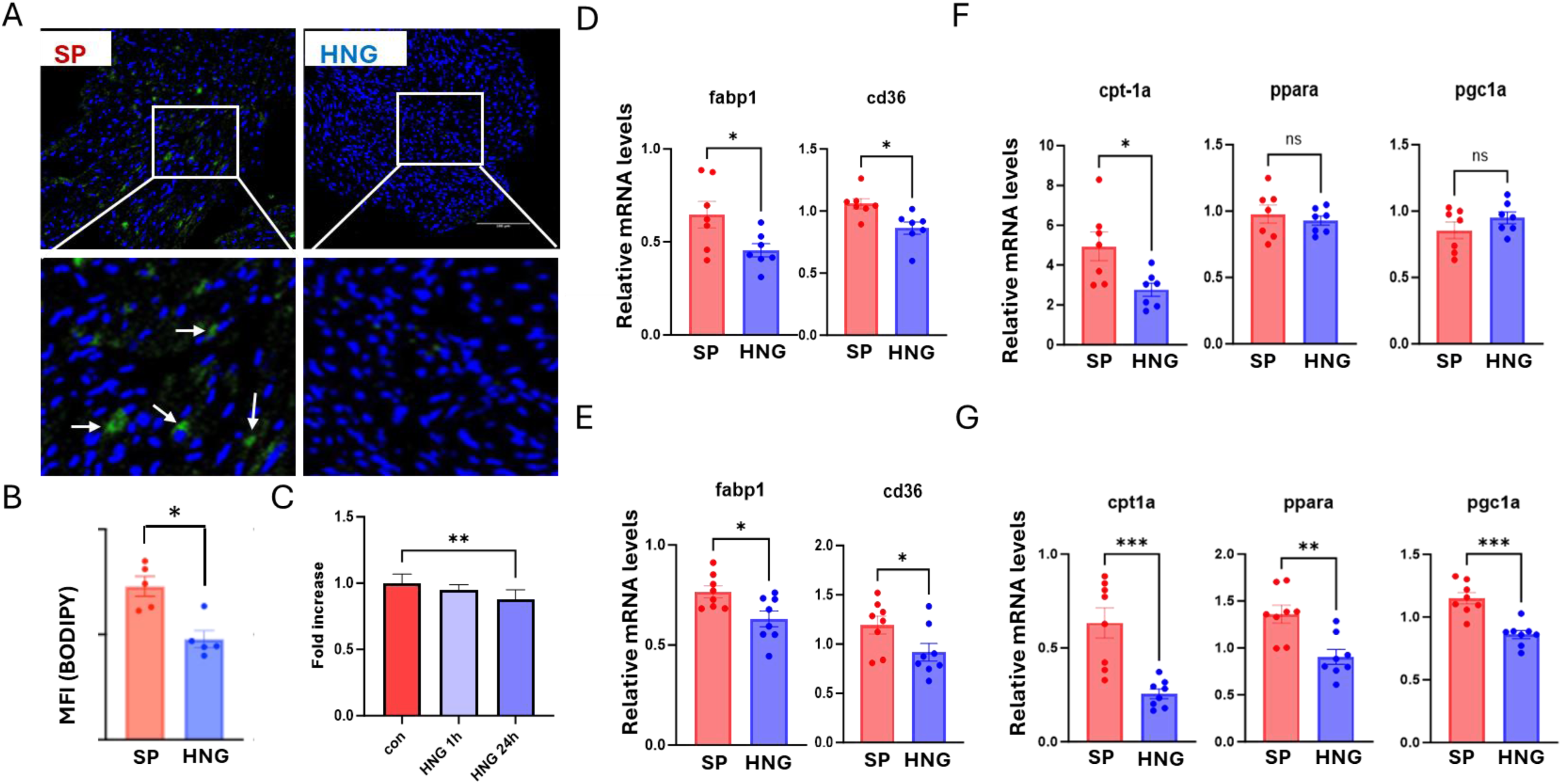
HNG treatment decreases fatty acid uptake and metabolism. A and B: Representative images (A) and quantification (B) for BODIPY labeled free fatty acid uptake in the heart in young female mice following MI and SP or HNG treatment; C: fatty acid uptake in H9C2 cells *in vitro* with the treatment of HNG for 1 hour or overnight; D and E: relative mRNA expression levels of *Fabp1 and Cd36* in both young (D) and old (E) female mouse heart tissue following MI and SP or HNG treatment; F-G: Relative mRNA expression levels of fatty acid metabolism markers, including *Cpt1a, Ppar α and Pgc 1α* in young (F) and old (G) mouse heart tissue following MI and SP or HNG treatment. Scale bars represent 100 μm. Results are shown as mean ± SEM. * p<0.05 and ** p<0.01

### HNG promotes glucose metabolism in the heart

To further assess the role of HNG on energetic flux, we examined ATP levels in H9C2 cells treated with HNG in the presence of different glucose levels in the media. HNG treatment increased ATP production (Fig. 9A, p<0.01) at both 5 mM and 10 mM glucose concentrations, indicating a possible increase in glucose metabolism by HNG. Indeed, assaying glucose oxidation using ^14^C radiolabeled glucose demonstrated that HNG significantly increased glucose oxidation to CO_2_ in H9C2 cells (Fig. 9B, p< 0.001). Glucose uptake in H9C2 cells increased following HNG treatment, with the increase in uptake seen as early as 15-min and steadily increasing up to 60 mins (Fig. 9C, p<0.001). Consistent with increased glucose uptake, Seahorse extracellular flux analysis demonstrated increased oxygen consumption rate in H9C2 cells with HNG treatment (Fig. 9D, p<0.05). To further dissect the effects of HNG on glucose metabolism, we performed glucose flux studies using ^13^Cstable isotope-labeled glucose. Consistent with the glucose uptake assay, there is an increase in uptake of ^13^C labeled glucose in the cells treated with HNG (Fig. 9E, p<0.01). Detailed analysis of metabolites showed that HNG treatment reduced lactate levels (p<0.05) but increased metabolites along the TCA cycles, including alpha-ketoglutarate (α-KG), succinate, malate, fumarate and aspartate (Fig. 9E, p<0.05), suggesting that HNG treatment increases flux through the TCA cycle in the heart. To further investigate whether this increased TCA cycle by HNG mediates the ATP production, we blocked the TCA cycle using aminooxyacetic acid (AOA), a known TCA cycle inhibitor. HNG induced increase in ATP production in H9C2 cells was completely abolished by treatment with 2mM of AOA (Fig. 9F, p<0.001). Taken together, HNG increases ATP production through increases in glucose uptake and oxidation in the cardiomyocytes.

**Fig. 9:**
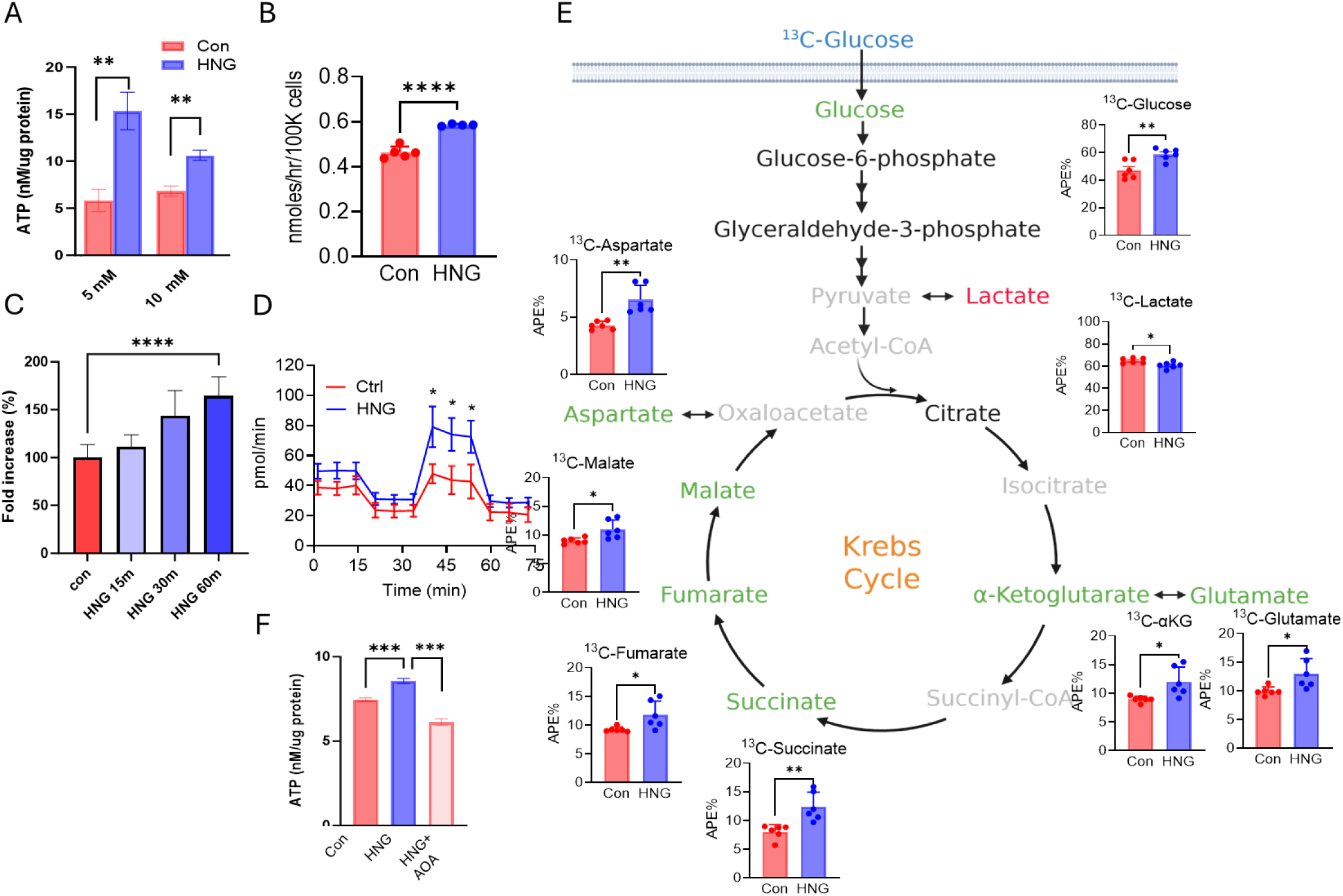
HNG treatment increases glucose metabolism. A: ATP production in H9C2 cells treated with 1 μM HNG under 0, 5 and 10mM glucose in the media; B: ^14^C-glucose oxidation rate in H9C2 cells treated with 1μM HNG; C: Glucose uptake in H9C2 cells treated with 1μM HNG for 15, 30 and 60 min; D: Oxygen consumption rate (OCR) in H9C2 cells treated with 1μM HNG using Seahorse; E: Glucose flux in H9C2 cells treated with 1μM HNG using ^13^C labeled glucose, shown as the total atom percent excess (APE%) for all 13C-labeled isotopologues; F: ATP production in H9C2 cells treated with 1μM HNG or HNG+AOA. Results are shown as mean ± SEM. * p<0.05, ** p<0.01 and *** p<0.001

### HNG increases GLUT4 translocation through VAMP7

To understand how HNG increases glucose uptake, we performed a GLUT4 translocation assay. Using both confocal microscopy (Fig. 10A) and flowcytometry (Fig. 10B, p<0.05), we show that HNG increases GLUT4 translocation to the cell surface. We then performed a proteomic study to identify potential HN interacting proteins by transfecting a GFP-HN fusion construct into H9C2 cells and pulling down HN using GFP antibody. This identified endocytosis and vesicle trafficking proteins including VAMP7, Rab7a, Gdi2 and Vps29 (Supp Table 1) as potential binding partners for HN. A Co-IP study confirmed that HN interacts with VAMP7 (Fig 10C) but not others (data not shown). To examine whether VAMP7 is the key mediator of HNG’s effect on glucose metabolism, we knocked down VAMP7 in HL-1 cells (Fig 10D, p<0.01) and showed that knockdown of VAMP7 abolishes HNG’s effect on GLUT4 translocation (Fig. 10E and F, p<0.05), and HNG-induced ATP production in cardiomyocytes (Fig. 10 G, p<0.05). Together, these data suggest that HNG mediates GLUT4 translocation to the cell surface through VAMP7.

**Fig. 10:**
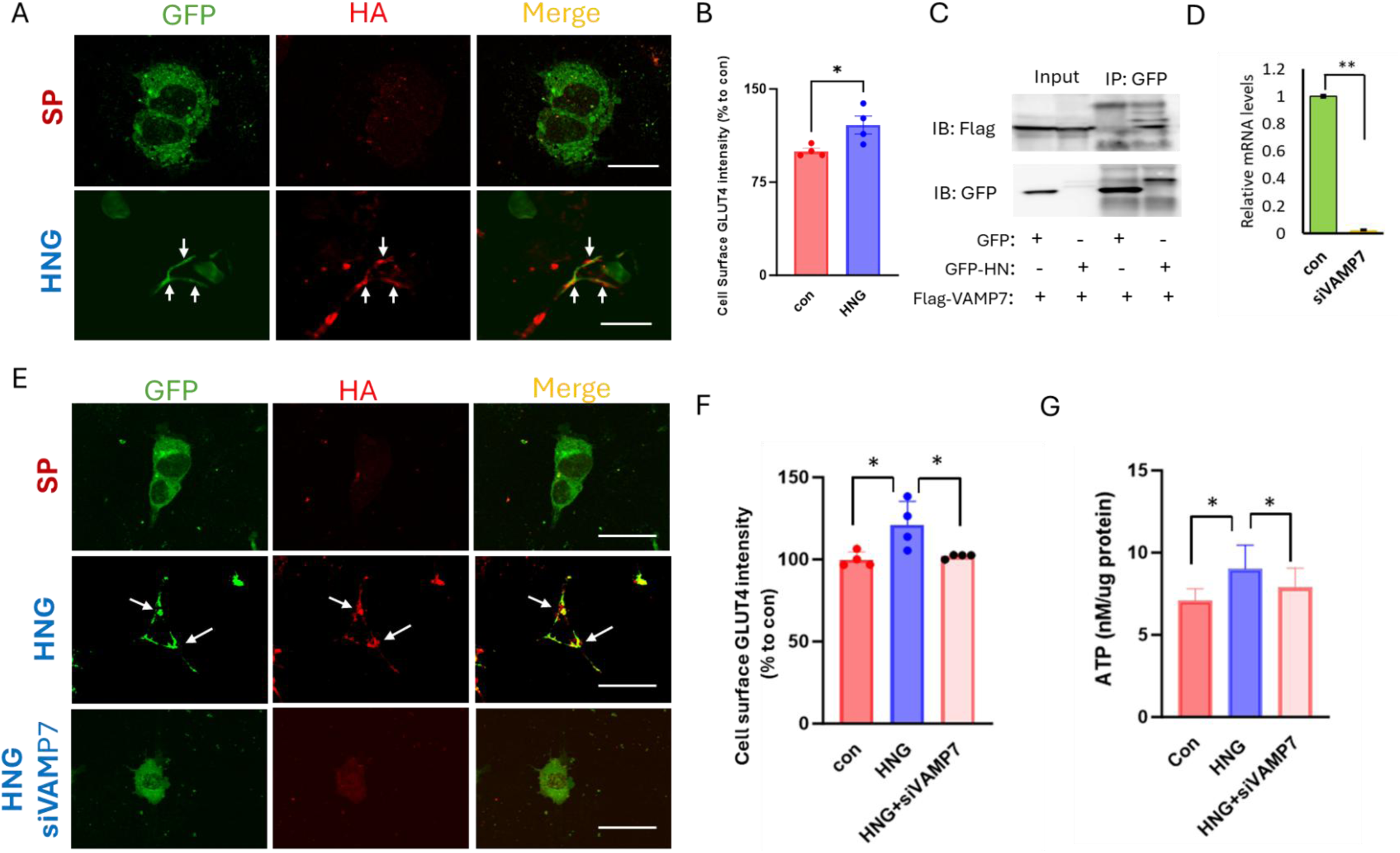
Effects of HNG on GLUT 4 translocation and glucose metabolism are VAMP 7 dependent. A and B: HNG-induced increase in cell surface HA-GLUT4-GFP was detected with anti-HA monoclonal Ab and Alexa Fluor 596-labeled goat anti-mouse IgG in non-permeabilized HL-1 cells using confocal microscopy (A) and flow cytometry (B); C: Interaction of VAMP7 and HN detected by co-IP followed by immunoblotting; D: expression levels of vamp7 in control or siVAMP7 knockdown HL-1 cells; E and F: Cell surface HA-GLUT4-GFP after siVAMP7 treatment by confocal microscopy (E) and flow cytometry (F); G: ATP production in HNG treated HL-1 cells with or without VAMP7 knockdown. Scale bars represent 50 μm. Results are shown as mean ± SEM. * p<0.05

## DISCUSSION

Our study demonstrates the cardioprotective effects of humanin analog HNG in the setting of chronic MI in both young and old mice. HNG treatment attenuated adverse ventricular remodeling, improved cardiac function, and induced a metabolic shift away from FAO, and towards glucose utilization, resulting in enhanced ATP production and cardiomyocyte survival. Mechanistically, we identify an interaction between HNG and VAMP7 that mediates GLUT4 translocation, promotes glucose uptake and oxidation, and increases myocardial ATP generation.

We previously showed that endogenous cardiac HN levels increase following myocardial ischemia (11), whereas circulating HN levels decline with advancing age in both mice and humans, implicating HN deficiency in age-related cardiovascular pathology (18). In murine MI/R models, a single dose of HNG administered either before ischemia or at reperfusion significantly reduced infarct size and preserved left ventricular function (19). Translational potential was demonstrated in a large-animal porcine MI/R model, in which HNG administration at reperfusion reduced myocardial infarct size (12). Independent studies by others showed that HNG limits infarct development and improves cardiac performance by attenuating mitochondrial dysfunction and oxidative stress during I/R (20, 21). Whereas these studies demonstrated benefit with acute HNG exposure in the short term, the present work is the first to establish that sustained HNG treatment favorably modulates cardiac remodeling and function in the context of chronic ischemia. These findings are also consistent with reports of reduced fibrosis in the heart following long-term HNG administration in aging rodent models (9) without an ischemic insult. Importantly, our chronic ischemia model isolates adaptive metabolic and remodeling responses in the absence of reperfusion injury, thereby enabling direct assessment of the effects of HNG on these processes. In addition to attenuating myocardial fibrosis, HNG treatment reduced the levels of expression of many oxidative stress response genes in the hearts of young mice, reflecting a lower oxidative stress in mice treated with HNG. Although a similar trend was observed in aged animals, this did not achieve statistical significance, potentially reflecting a higher baseline oxidative burden with aging—an issue that warrants further investigation.

Myocardial metabolomic analyses revealed robust metabolic reprogramming in response to HNG. HNG-treated hearts exhibited significantly reduced levels of multiple fatty acids, accompanied by downregulation of key FAO regulators, including *PPARα, PGC*-*1α,* and *CPT1,* in both young and old mice, with more pronounced effects in the aged cohort. These findings indicate that reduced myocardial fatty acid content did not result from enhanced fatty acid oxidation. Indeed, a downregulation of PPARα has been shown to prevent lipotoxicity in ischemic cardiomyopathy (22). Because impaired β-oxidation capacity can lead to accumulation of toxic lipid intermediates and lipotoxic cardiomyocyte injury (23), we also examined fatty acid uptake. HNG treatment decreased *in vivo* BODIPY labelled palmitate uptake, and reduced expression of the fatty acid transporters *Cd36* and *Fabp* indicating diminished myocardial fatty acid uptake. These results align with prior studies demonstrating that limiting fatty acid uptake or CD36 activity protects against lipotoxicity, oxidative stress, and ischemic cardiomyopathy (24–28).

Although ischemic myocardium typically suppresses FAO as an adaptive response to hypoxia, preservation of energetic homeostasis requires compensatory utilization of alternative substrates. Chronic ischemia is associated with a reversion to a “fetal” metabolic phenotype characterized by enhanced glucose dependence. Our data demonstrates that HNG increases glucose uptake and promotes this metabolic shift in a favorable manner by enhancing glucose oxidation rather than glycolysis alone. Metabolic flux analyses revealed increased TCA cycle activity, improved coupling of glycolysis to mitochondrial oxidation, reduced lactate accumulation, and increased ATP production. Inhibition of the TCA cycle abolished HNG-induced ATP generation, underscoring the potential role of increased glucose oxidation in the HNG-mediated cardioprotection. Time-course studies showed that the increase in glucose oxidation and ATP production preceded reductions in FAO (Randle’s effect), indicating that enhanced glucose oxidation is the primary driver of metabolic reprogramming. These findings are concordant with prior studies demonstrating that stimulation of glucose oxidation using dichloroacetate improves energetic efficiency, reduces oxidative stress, and enhances cardiac function under ischemic conditions (29). GLUT4, the predominant glucose transporter in the heart, is a key mediator of ischemia-induced glucose uptake (30). Heather et al. demonstrated an ischemia-associated 32% reduction in sarcolemmal FAT/CD36 and a 90% increase in sarcolemmal GLUT4 following ischemia (31). Consistent with this, *Glut4⁻^/^⁻* hearts exhibit profound and irreversible systolic and diastolic dysfunction following I/R, while cardiac-specific deletion of pyruvate dehydrogenase, the key enzyme that links glycolysis to the TCA cycle, impairs glucose oxidation, increases infarct size, and induces diastolic dysfunction (32). Nevertheless, myocardial substrate utilization does not reflect a simple binary switch between glucose and FAO. Clinical conditions such as insulin resistance and obesity substantially alter coordinated substrate metabolism and may contribute to the progression of heart failure.

Given the central role of GLUT4 in myocardial glucose uptake during ischemia, we next investigated the mechanism by which HNG promotes GLUT4 translocation. Using a proteomics-based approach, we identified VAMP7 as a binding partner of HNG. Prior studies have established VAMP7 as a key regulator of GLUT4 trafficking to and from the plasma membrane (33, 34). We demonstrate that the HNG–VAMP7 interaction is required for GLUT4 translocation under identical glucose and insulin conditions and that VAMP7 knockdown abolishes HNG-induced GLUT4 membrane localization and ATP production. These findings identify HNG–VAMP7–dependent GLUT4 trafficking as a critical mechanism linking metabolic reprogramming to cardioprotection during chronic ischemia.

All chronic ischemia studies were performed in female mice, as male control mice exhibited high post-ligation mortality (S Fig 1) consistent with prior reports. In contrast, prior murine acute ischemia studies were conducted in males, whereas porcine acute I/R studies were performed in females (11, 12). Thus, HNG appears to confer cardioprotection across sexes; however, future studies will be required to rigorously define sex-specific mechanisms and therapeutic responses to HNG in the setting of chronic ischemia. Finally, HN is endogenously produced and has been well tolerated in preclinical studies with treatment durations of up to 14 months (9, 18), underscoring the translational potential of HN and its analogs for myocardial infarction–induced heart failure.

### Significance

Collectively, these findings identify Humanin as a novel endogenous modulator of myocardial metabolism that confers cardioprotection in the setting of chronic ischemia. By coordinating suppression of fatty acid uptake with enhancement of glucose oxidation through a VAMP7–GLUT4–dependent mechanism, HNG promotes energetically efficient substrate utilization that preserves myocardial ATP levels, limits lipotoxic and oxidative injury, and attenuates adverse remodeling. Importantly, these effects are observed in both young and aged hearts, addressing a critical unmet need for therapies that remain effective across the aging spectrum. Given the endogenous origin, favorable safety profile, and sustained efficacy of HN analogs, these data support targeting HNG–mediated metabolic reprogramming as a promising therapeutic strategy for ischemic heart disease and post–myocardial infarction heart failure.

---

**Fig. S1:**
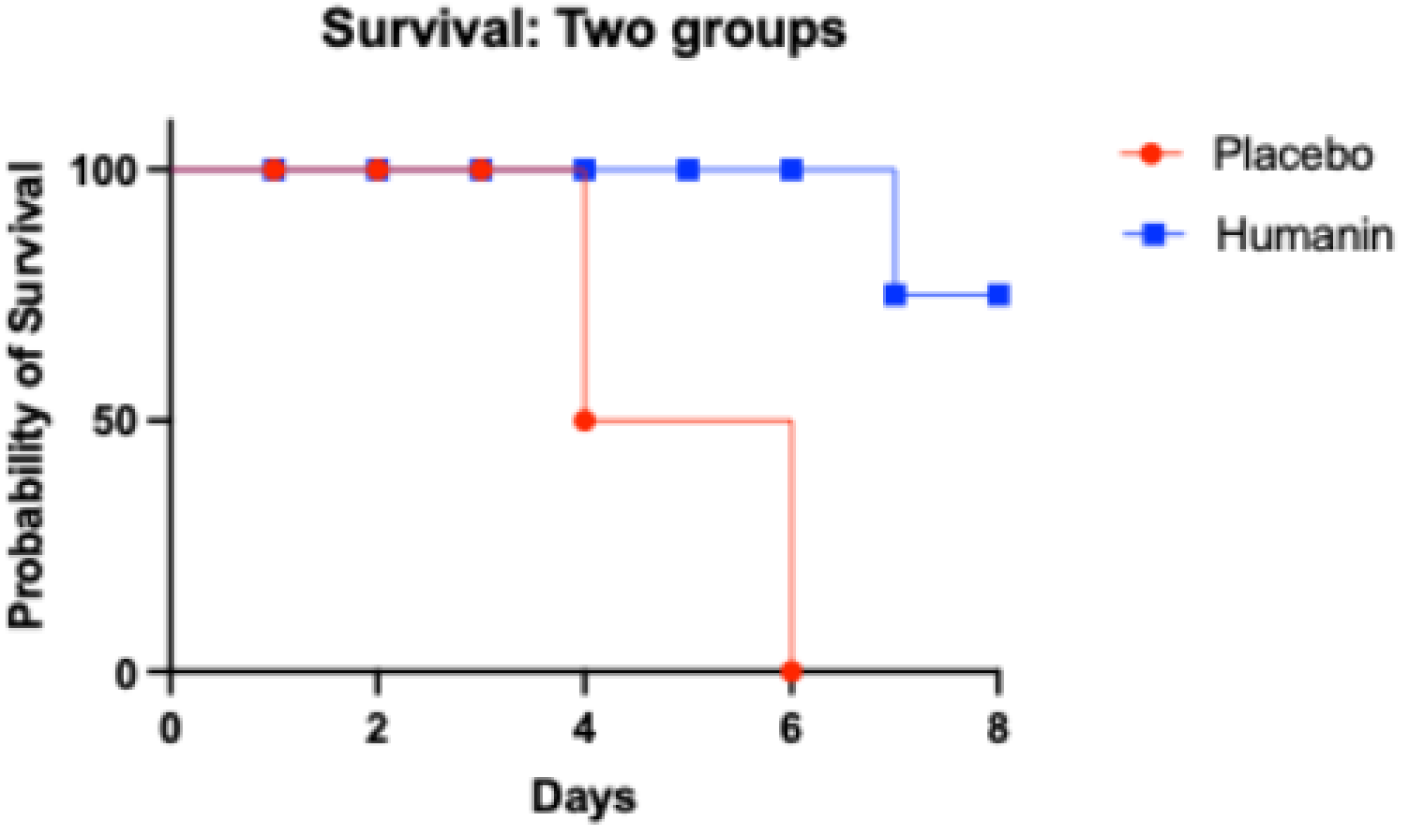
HNG increases survival rate in young male mice following chronic MI.

**Supp Table 1:**
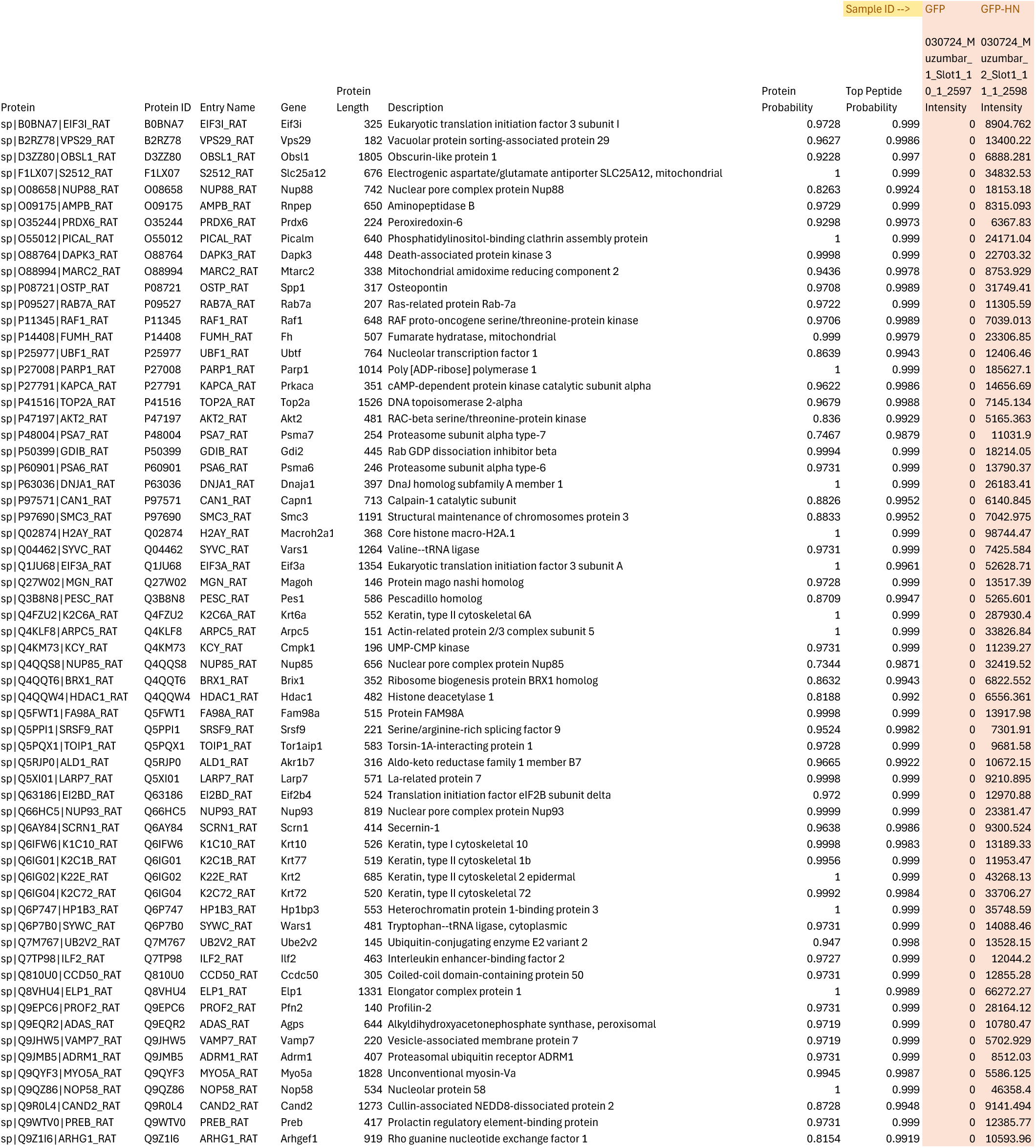
Proteomics.

## References

1. B. Bozkurt et al., Heart Failure Epidemiology and Outcomes Statistics: A Report of the Heart Failure Society of America. J Card Fail 29, 1412–1451 (2023).

2. T. Yan et al., Burden, Trends, and Inequalities of Heart Failure Globally, 1990 to 2019: A Secondary Analysis Based on the Global Burden of Disease 2019 Study. Journal of the American Heart Association 12, e027852 (2023).

3. C. W. Tsao et al., Heart Disease and Stroke Statistics-2023 Update: A Report From the American Heart Association. Circulation 147, e93–e621 (2023).

4. A. Kochar et al., Long-Term Mortality of Older Patients With Acute Myocardial Infarction Treated in US Clinical Practice. Journal of the American Heart Association 7 (2018).

5. D. M. Yellon, D. J. Hausenloy, Myocardial reperfusion injury. N Engl J Med 357, 1121–1135 (2007).

6. S. Li, B. Xie, W. Zhang, T. Li, Dichloroacetate ameliorates myocardial ischemia-reperfusion injury via regulating autophagy and glucose homeostasis. Arch Med Sci 19, 420–429 (2023).

7. C. J. Holubarsch et al., A double-blind randomized multicentre clinical trial to evaluate the efficacy and safety of two doses of etomoxir in comparison with placebo in patients with moderate congestive heart failure: the ERGO (etomoxir for the recovery of glucose oxidation) study. Clin Sci (Lond) 113, 205–212 (2007).

8. Y. Hashimoto et al., A rescue factor abolishing neuronal cell death by a wide spectrum of familial Alzheimer’s disease genes and Abeta. Proceedings of the National Academy of Sciences of the United States of America 98, 6336–6341 (2001).

9. Q. Qin et al., Chronic treatment with the mitochondrial peptide humanin prevents age-related myocardial fibrosis in mice. American journal of physiology. Heart and circulatory physiology 315, H1127–H1136 (2018).

10. Z. Gong, E. Goetzman, R. H. Muzumdar, Cardio-protective role of Humanin in myocardial ischemia-reperfusion. Biochim Biophys Acta Gen Subj 1866, 130066 (2022).

11. R. H. Muzumdar et al., Humanin: a novel central regulator of peripheral insulin action. PLoS One 4, e6334 (2009).

12. T. E. Sharp, 3rd et al., Efficacy of a Novel Mitochondrial-Derived Peptide in a Porcine Model of Myocardial Ischemia/Reperfusion Injury. JACC. Basic to translational science 5, 699–714 (2020).

13. E. Johny, P. Dutta, Left coronary artery ligation: a Surgical Murine Model of myocardial infarction. Journal of visualized experiments: JoVE, 10.3791/64387 (2022).

14. A. Zeigerer et al., GLUT4 retention in adipocytes requires two intracellular insulin-regulated transport steps. Mol Biol Cell 13, 2421–2435 (2002).

15. L. E. Klein, L. Cui, Z. Gong, K. Su, R. Muzumdar, A humanin analog decreases oxidative stress and preserves mitochondrial integrity in cardiac myoblasts. Biochem Biophys Res Commun 10.1016/j.bbrc.2013.08.055 (2013).

16. Z. Gong et al., Central effects of humanin on hepatic triglyceride secretion. Am J Physiol Endocrinol Metab 309, E283–292 (2015).

17. R. Kuliawat et al., Potent humanin analog increases glucose-stimulated insulin secretion through enhanced metabolism in the beta cell. FASEB J 27, 4890–4898 (2013).

18. L. J. Cobb et al., Naturally occurring mitochondrial-derived peptides are age-dependent regulators of apoptosis, insulin sensitivity, and inflammatory markers. Aging 8, 796–809 (2016).

19. R. H. Muzumdar et al., Acute humanin therapy attenuates myocardial ischemia and reperfusion injury in mice. Arterioscler Thromb Vasc Biol 30, 1940–1948 (2010).

20. S. Thummasorn et al., Humanin exerts cardioprotection against cardiac ischemia/reperfusion injury through attenuation of mitochondrial dysfunction. Cardiovascular therapeutics 34, 404–414 (2016).

21. S. Thummasorn, K. Shinlapawittayatorn, S. C. Chattipakorn, N. Chattipakorn, High-dose Humanin analogue applied during ischemia exerts cardioprotection against ischemia/reperfusion injury by reducing mitochondrial dysfunction. Cardiovascular therapeutics 35 (2017).

22. O. Dewald et al., Downregulation of peroxisome proliferator-activated receptor-alpha gene expression in a mouse model of ischemic cardiomyopathy is dependent on reactive oxygen species and prevents lipotoxicity. Circulation 112, 407–415 (2005).

23. K. D’Souza, C. Nzirorera, P. C. Kienesberger, Lipid metabolism and signaling in cardiac lipotoxicity. Biochimica et biophysica acta 1861, 1513–1524 (2016).

24. Y. Angin et al., CD36 inhibition prevents lipid accumulation and contractile dysfunction in rat cardiomyocytes. Biochem J 448, 43–53 (2012).

25. D. P. Koonen, J. F. Glatz, A. Bonen, J. J. Luiken, Long-chain fatty acid uptake and FAT/CD36 translocation in heart and skeletal muscle. Biochimica et biophysica acta 1736, 163–180 (2005).

26. J. Yang et al., CD36 deficiency rescues lipotoxic cardiomyopathy. Circulation research 100, 1208–1217 (2007).

27. J. F. Glatz, J. J. Luiken, A. Bonen, Membrane fatty acid transporters as regulators of lipid metabolism: implications for metabolic disease. Physiological reviews 90, 367–417 (2010).

28. J. C. Perman et al., The VLDL receptor promotes lipotoxicity and increases mortality in mice following an acute myocardial infarction. The Journal of clinical investigation 121, 2625–2640 (2011).

29. G. D. Lopaschuk, Q. G. Karwi, R. Tian, A. R. Wende, E. D. Abel, Cardiac Energy Metabolism in Heart Failure. Circulation research 128, 1487–1513 (2021).

30. R. Tian, E. D. Abel, Responses of GLUT4-deficient hearts to ischemia underscore the importance of glycolysis. Circulation 103, 2961–2966 (2001).

31. L. C. Heather et al., Differential translocation of the fatty acid transporter, FAT/CD36, and the glucose transporter, GLUT4, coordinates changes in cardiac substrate metabolism during ischemia and reperfusion. Circulation. Heart failure 6, 1058–1066 (2013).

32. K. Gopal et al., Cardiac-Specific Deletion of Pyruvate Dehydrogenase Impairs Glucose Oxidation Rates and Induces Diastolic Dysfunction. Frontiers in cardiovascular medicine 5, 17 (2018).

33. D. Williams, J. E. Pessin, Mapping of R-SNARE function at distinct intracellular GLUT4 trafficking steps in adipocytes. The Journal of cell biology 180, 375–387 (2008).

34. R. W. Schwenk et al., Requirement for distinct vesicle-associated membrane proteins in insulin- and AMP-activated protein kinase (AMPK)-induced translocation of GLUT4 and CD36 in cultured cardiomyocytes. Diabetologia 53, 2209–2219 (2010).

